# De Novo-Designed Alpha-Helical Proteins Enable Ligand-Directed Spectral Tuning

**DOI:** 10.64898/2026.11.26.769376

**Authors:** Jiale Liu, Yan Xiong, Wei Zhu, Chengyang Li, Zijie Cheng, Zheng Guo, Yanyan Wang, Jiaxuan Chen, Yueqin Zheng, Minjie Sun, Luhua Lai

## Abstract

Canonical green fluorescent proteins typically regulate chromophore photophysics through a rigid beta-barrel environment, while spectral tuning often relies on protein engineering or directed evolution. Here, we design a compact alpha-helical protein pocket that activates a GFP-chromophore-derived fluorogen and accommodates a series of chemically modified ligands. A co-crystal structure reveals shared anchoring interactions for fluorogen activation and contacts that enable ligand-dependent spectral tuning. Guided by this structure, ligand substitution shifts emission across the red-to-green range while retaining the same protein scaffold. Quantitative binding and photophysical analyses establish the structural and chemical determinants of affinity, fluorescence activation, and spectral tuning, and the resulting fluorogens function in living cells. Finally, structure-guided substitutions repurpose the pocket toward a distinct drug ligand. These results establish an alpha-helical route to programmable fluorogen activation in which optical properties can be tuned through ligand design rather than repeated protein evolution.

## Introduction

The fluorescence of GFP-derived chromophores is strongly dependent on the molecular environment surrounding the chromophore [1]. In the canonical green fluorescent protein, the chromophore is generated and immobilized within a rigid beta-barrel, where a precisely organized network of residues restricts internal motion and modulates local elec-trostatics, protonation, and nonradiative relaxation [2]. The GFP beta-barrel therefore provides more than a structural enclosure: it creates a photophysical microenvironment that converts a weakly fluorescent chromophore into a bright fluorophore.

Fluorogen activation is not, however, limited to the GFP beta-barrel architecture. Recent studies have shown that compact alpha-helical protein scaffolds can activate other classes of fluorogens, such as HBC-based dyes [3] and rhodamine-based dyes [4, 5]. These studies establish the feasibility of using helical protein environments to restrict fluorogen motion and generate fluorescence turn-on. In parallel, FAST-family chemogenetic tags have been engineered to bind chemically related fluorogens and generate different spectral outputs from a common protein scaffold [6]. More recently, de novo-designed pan-rhodamine binders have been developed to accommodate diverse rhodamine fluorophores [5]. However, GFP-chromophore-derived fluorogens differ from these dyes in their molecular geometry, conjugation pattern, and photophysical requirements. Whether such ligands can be recognized and activated within a de novo alpha-helical pocket, while allowing ligand-dependent spectral tuning, remains unclear.

The relevant unresolved question is therefore not whether a protein environmen-t can activate a fluorogen in general, but whether a alpha-helical microenvironment can reproduce the specific structural and chemical constraints required for GFP-chromophore-derived fluorescence. A related question is whether one such pocket can accommodate a series of chemically modified GFP-chromophore derivatives while preserving binding and fluorogenicity. In established fluorescent-protein systems, spectral properties are commonly tuned by modifying the protein scaffold through mutagenesis or directed evolution. A protein-ligand system in which spectral output can instead be adjusted through ligand chemistry would provide a complementary route to fluorogen engineering, while avoiding the need to redesign a protein scaffold for each optical color, and helping us understand protein–chromophore recognition and fluorescence activation beyond the GFP beta-barrel.

Alpha-helical bundles provide a distinct framework for addressing these questions. Their geometry is compact, modular, and amenable to de novo construction [7–10], allowing the organization of binding pockets with defined spatial relationships between hydrophobic, aromatic, and polar side chains [11–15]. For fluorogenic ligands, however, affinity alone is insufficient. The pocket must simultaneously anchor the ligand, restrict chromophore motion, and create a local environment compatible with fluorescence activation [16–18]. A successful design must therefore couple molecular recognition to the construction of a functional photophysical microenvironment.

Here, we establish a de novo alpha-helical fluorogenic pocket for a GFP-chromophore-derived ligand. Using a computationally guided design strategy, we identified a compact helical protein binder and determined its ligand-bound structure at 1.6 Å resolution. The structure reveals how the protein recognizes distinct regions of the ligand and organizes the chromophore within the pocket. Mutagenesis further showed that a single hydrogen-bonding interaction couples ligand affinity to chromophore photophysics: disrupting this contact shifted the excitation maximum by approximately 40 nm while weakening binding by approximately 38-fold. Guided by these structural features, we designed a series of GFP-chromophore derivatives that bind the same protein scaffold and shift emission across the red-to-green spectral range without changing the protein sequence. Quantitative binding, photophysical, and cellular analyses define the contributions of ligand structure to affinity, fluorescence activation, and spectral output. Finally, structure-guided substitutions at three pocket positions redirect the protein toward a chemically distinct drug ligand. Together, these results establish a GFP-chromophore-based alpha-helical fluorogenic system in which a structurally defined protein microenvironment supports ligand-directed spectral tuning and programmable ligand recognition.

## Results

### Protein-directed versus ligand-directed tuning of chromophore fluorescence

In canonical GFP systems, chromophore fluorescence is tuned primarily through the protein scaffold [19–21]. Mutations surrounding the chromophore alter its conformational freedom, electrostatic environment, and protonation state, thereby producing distinct spectral properties. This protein-directed mode of tuning has enabled the development of fluorescent proteins with diverse colors, but each new optical output generally requires modification of the protein sequence or selection of a new protein variant (Fig 1A).

**Figure 1:**
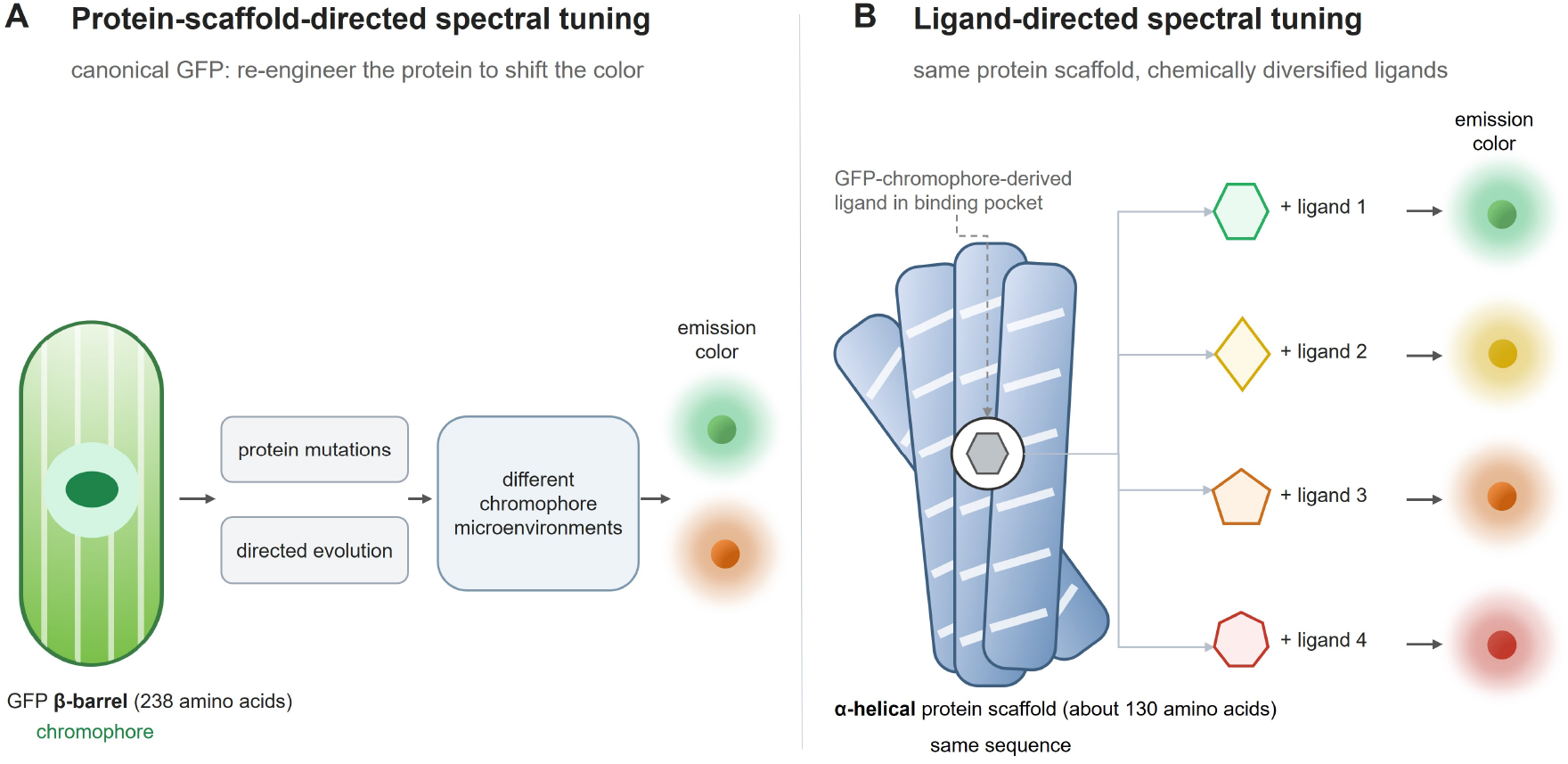
Protein-directed versus ligand-directed tuning of chromophore fluorescence. **A** In canonical GFP systems, spectral properties are tuned by modifying residues surrounding the chromophore, thereby changing the protein microenvironment. **B** In this work, a de novo alpha-helical protein scaffold is held constant while a series of GFP-chromophore-derived ligands is used to tune the emission wavelength.

We asked whether the same conceptual function could be implemented through ligand chemistry in a de novo protein environment. Specifically, we sought to establish an alpha-helical pocket that could activate a GFP-chromophore-derived fluorogen and accommodate chemically modified derivatives without redesigning the protein scaffold for each ligand. In this design, the protein provides a conserved fluorogenic microenvironment, whereas ligand substitution provides the variable element that controls emission wavelength (Fig 1B).

To test this concept, we first needed to establish an alpha-helical protein capable of binding and activating a GFP-chromophore-derived fluorogen. We therefore applied ChameleonBinder to the design of DFAME-binding proteins and evaluated the resulting designs experimentally. Briefly, the workflow of ChameleonBinder combines scaffold library construction [10, 22, 23], global ligand docking with pseudo-Vals [24–26], ligand-aware sequence design [27], and complex structure prediction [28] to identify self-consistent alpha-helical binding pockets. The detailed scaffold library, docking, sequence design, structure prediction, and filtering procedures are described in the Supplementary Information.

### Efficient computational design of DFAME-activating proteins with one-round experimental validation

DFAME, a red-emitting GFP-chromophore-derived fluorogen [29], has four rotatable bonds with molecular weight of 322.26 Da. To ensure its fluorescence emission after protein binding, we fixed two rotatable bonds to make it planar with two remaining bond rotatable. DFAME exhibits long-wavelength fluorescent emission and negligible autofluores-cence, suggesting the potential for developing protein-based fluorescence-imaging probes with reduced background interference and improved suitability for deep-tissue imaging. After three rounds of design and selection using ChameleonBinder (see Methods for details), 7 sequences were selected for experimental testing.

As shown in Fig 2A and B, the designed protein DFA-5 forms hydrogen bonds with nearby residues and exhibits high pLDDT confidence scores (*≥* 95) in both the apo and holo states. The holo structures have slightly higher pLDDT scores than the apo structures, suggesting a more stable conformation. Protein expression testing using *E.coli* showed that all of the designed proteins are soluble (Fig S6). After cell lysis, two of the supernanant showed significant fluorescent emission. We further purified these two proteins, and the elution volume from size exclusion chromatography (SEC) indicated that DFA-5 is homogenous, corresponding to a monomeric state (Fig 2C), while DFA-4 appeared as a mixture of monomers and dimers with more monomers (Fig S7).

**Figure 2:**
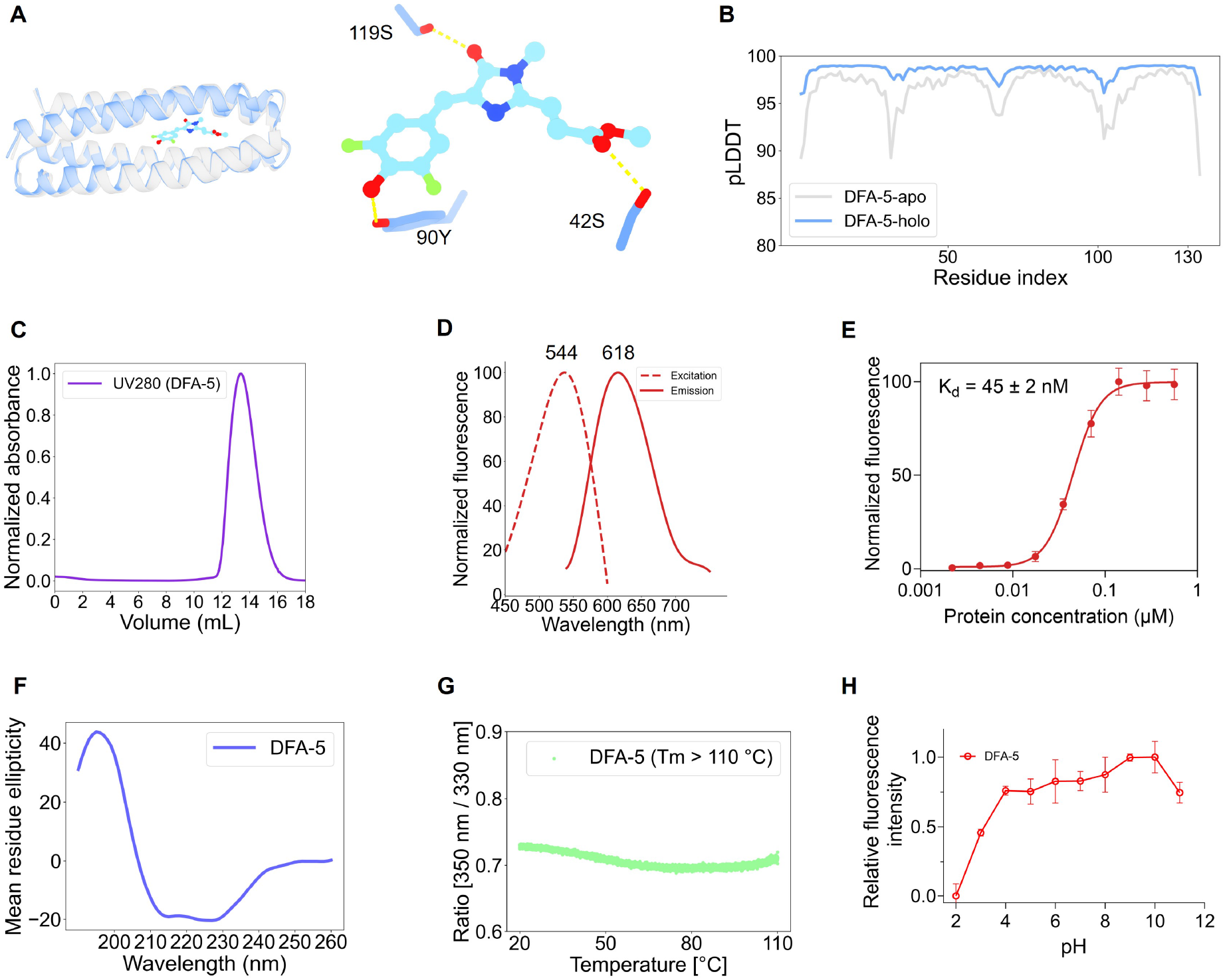
ChameleonBinder successfully designed DFAME-activating proteins. **A** and **B** show the designed protein DFA-5, including the predicted hydrogen bonds and high confidence scores. **C** to **H** present the experimental results of elution volume, circular dichroism spectra, spectral information of excitation and emission, binding affinity, melting temperatures, and fluorescence responses to pH changes, respectively.

We further analyzed the secondary structures using circular dichroism (CD), and DFA-5 displayed typical helical structural features (Fig 2F). Next, we conducted excitation and emission scans to determine the maximum values (Fig 2D) and measured the binding affinity based on the maximum emission when excited at its optimal excitation wavelength. The apparent *K_d_* value for DFA-5 binding to DFAME was found to be 45 nM (Fig 2E), which is three times lower than that of DFA-4 (Fig S7). DFAME-binding protein DFA-5 demonstrates a substantial Stokes shift of approximately 74 nm (Ex maxima at 544 nm and Em maxima at 618 nm), making it excellent candidate for minimizing interference from excitation signals. We further measured the melting temperatures (Tm) of DFA-5 (Fig 2G) and the fluorescence ratio changes (330 nm/350 nm) remained below 0.1 as the temperature increased to 110 *^◦^*C, indicating that DFA-5 is highly stable. We also assessed pH sensitivity and found that DFA-5 is stable across a broad pH range (4-11) (Fig 2H). Together, these results establish that a compact de novo alpha-helical protein can bind and activate DFAME, while exhibiting thermal and pH stability.

### Structural validation of the designed protein DFA-5

We successfully solved the crystal structure of DFA-5 in the presence of DFAME at high resolution 1.6 Å (Table S1, S2), which clearly show the electron density of DFAME and its nearby residues (Fig 3A). The conformational core of DFAME in the crystal structure is highly planar, where the dihedral torsion angles between the six and five-membered rings, the five-membered ring and the adjacent C=C double bond, the C=C double bond and carbonyl oxygen group (C=O) are 0.2°, 8.0°, and 0.7°, respectively. We also noticed that the predicted conformational core of DFAME using AF3 is similarly planar to the crystal conformation. However, the last two torsion angles above mentioned flipped around 180°, resulting in the position of carbonyl oxygen atom slightly changed (Fig 3B). We further checked the PDB and find that both our crystal structure and AF3 prediction structure are different to the three currently deposited aptamer-DFAME entries (PDB ID: 8EYU [30], 8FI0 and 8FI8 [31]). Structural alignment revealed that the designed protein model agrees well with the crystal structure where the RMSD of all C*α*, C*α* of residues in the ligand binding sites and C*α* of residues in the hydrophobic core are 0.67 Å, 0.50 Å, and 0.37 Å, respectively. Three well-designed hydrogen bonds are also presented in the crystal structures (Fig 3C). The hydroxyl in DFAME formed hydrogen bond with the phenolic hydroxyl of 90Y, the carbonyl oxygen in the five-membered ring in DFAME formed hydrogen bond with the hydroxyl group of 119S, and the carbonyl oxygen in DFAME formed hydrogen bond with the hydroxyl group of 42S.

**Figure 3:**
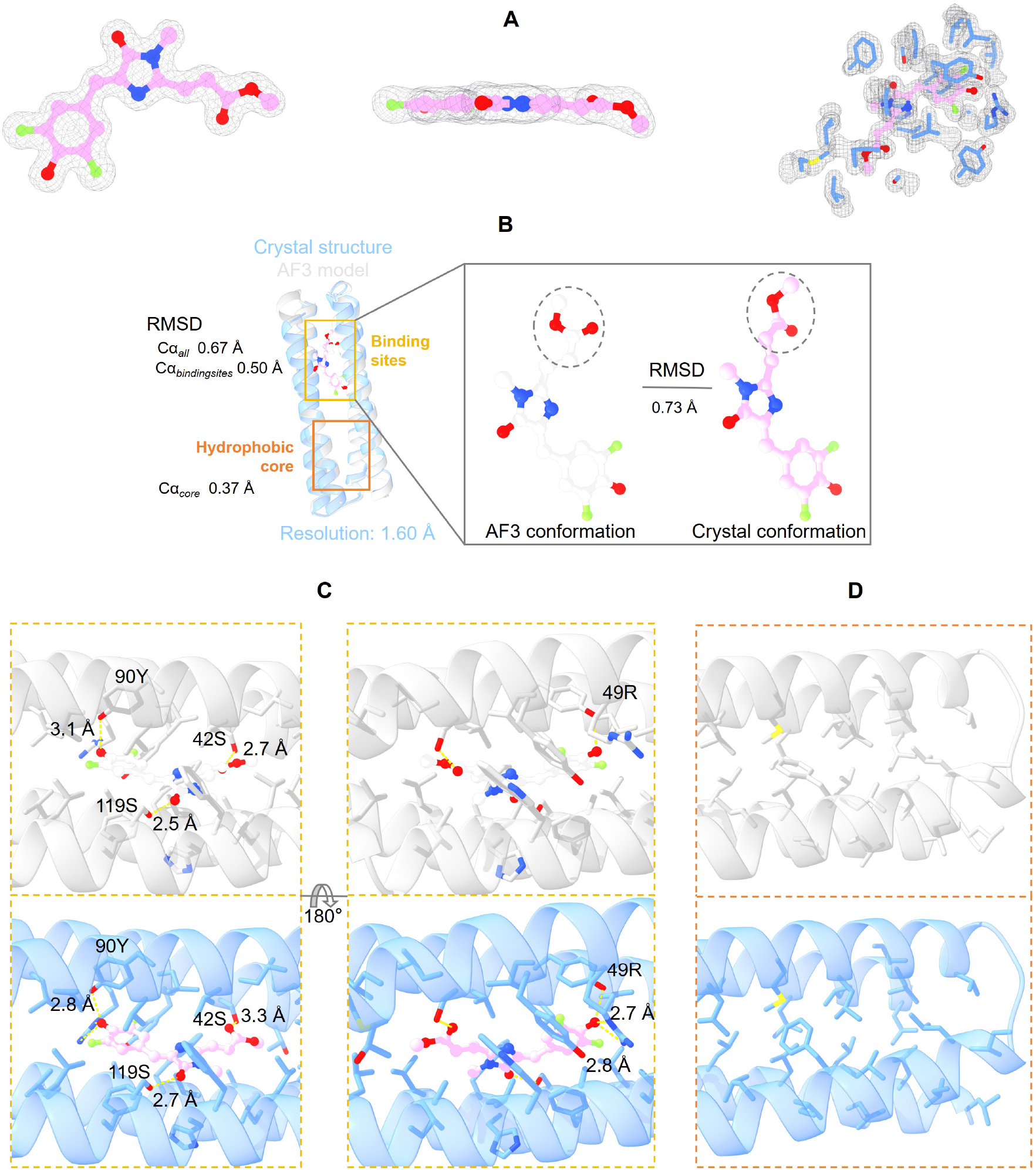
The solved co-crystal structure validates the designed model DFA-5. **A** 2mFo-DFc electron density map contoured at 1.0*σ* for the ligand DFAME and side chains of the surrounding residues. **B** Structural alignment between the AF3-predicted model and the solved crystal structure, with the enlarged part highlighting a flipped ligand conformation. **C** Detailed comparison of the binding site region. **D** Detailed comparison of the hydrophobic core.

We also noticed that there are two additional hydrogen bonds formed between the guanidinium group of 49R and the hydroxyl group in DFAME that are not appeared in the designed model due to the side-chain flexibility of 49R.

The conformations of hydrophobic core residues are close between the crystal structure and the designed model which is crucial to support the protein folding ability in the absence of ligand (Fig 3D). Together, these results show that the experimentally determined structure of DFA-5 closely matches the designed model.

### Analysis of fluorescence activation and microenvironment comparison between the designed proteins and natural fluorescent proteins

To investigate how specific residues in protein DFA-5 influence binding affinity and ability of fluorescence activation, we rationally mutated several key residues involved in forming buried hydrogen bonds with DFAME, including Y90F, S42A, S119A, and R49A. We then measured the fluorescence intensity using a consistent protein concentration (10 µM) and ligand concentration (500 nM) at the optimal excitation wavelength for these variants. The measurements were conducted at the maximum emission values. Notably, the fluorescence was almost completely abolished for the mutations S119A and R49A, while the mutant S42A displayed fluorescence intensity five times higher than that of DFA-5 (Fig 4A,B). Interestingly, the maximum excitation wavelength of the S42A mutant shifted substantially from 544 nm to 583 nm, and the maximum emission wavelength also shifted from 618 nm to 638 nm (Fig 4C). The apparent *K_d_* value for the S42A mutant was found to be 40 times higher than that of the original DFA-5 (Fig 4D). The S42 residue in DFA-5 is water-accessible (Fig 4E) and involved in hydrogen bonding with DFAME, as indicated by the crystal structures, allowing for rapid energy dissipation. Mutating S42 to Ala interrupts the hydrogen bonds, which may block the water-mediated energy transfer and enhance fluorescence emission.

**Figure 4:**
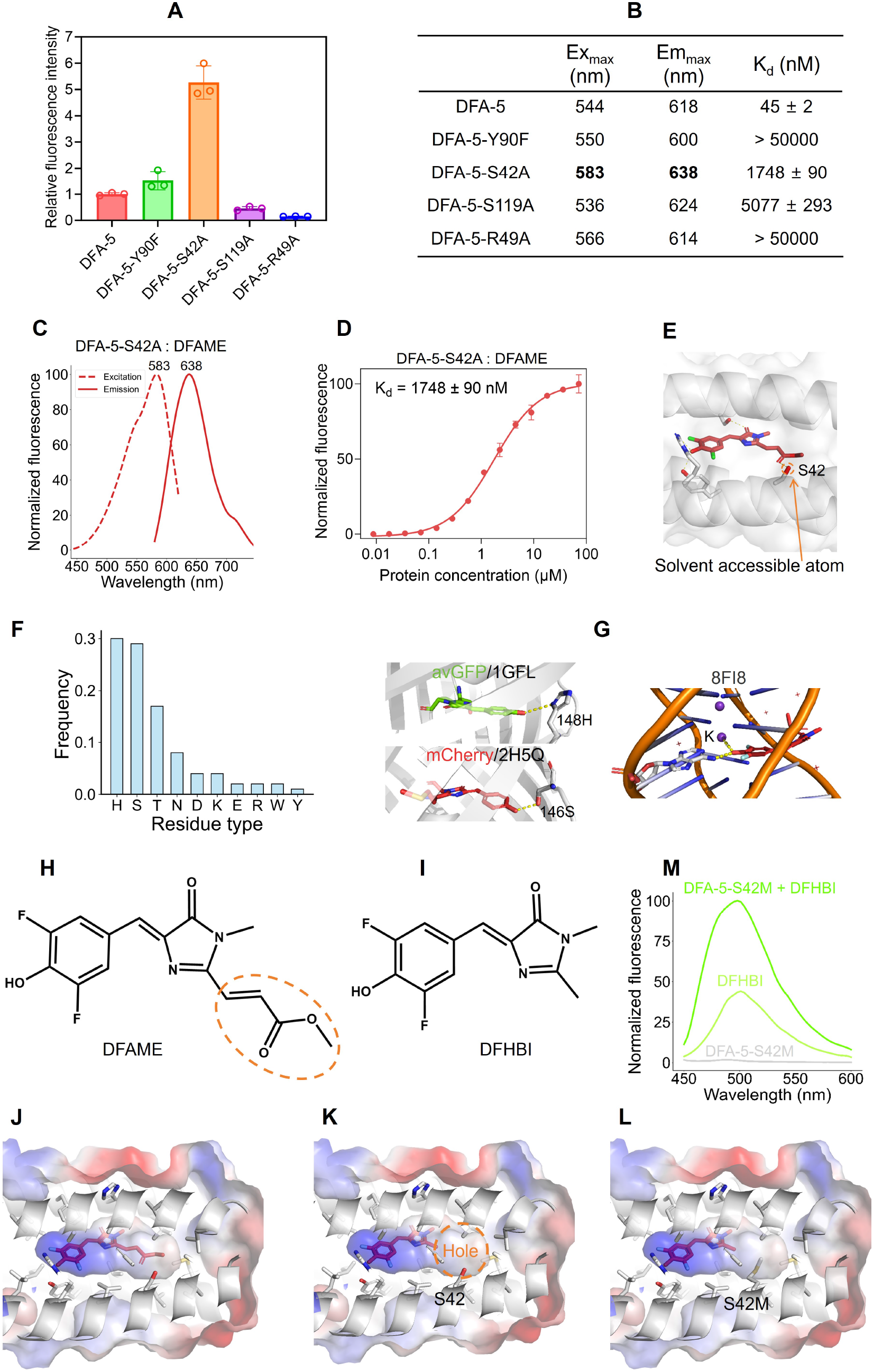
Microenvironment analysis reveals the mechanism of fluorescence activation. **A** Fluorescence intensity comparison between the designed protein DFA-5 and several variants. **B** Fluorescence properties and binding affinities. **C** The single-mutation variant DFA-5-S42A exhibits a larger red shift than DFA-5. **D** Binding affinity between DFA-5-S42A and DFAME. **E** Relationship between the increased fluorescence intensity and decreased binding affinity in DFA-5-S42A. **F** Statistical analysis of phenolic hydroxyl groups in native fluorescent proteins reveals distinct residue preferences compared with designed DFA-5. **G** Case studies showing different interaction patterns. **H** Molecular structures of DFAME and **I** DFHBI. **J,K** Replacement of DFAME with DFHBI leaves a cavity, resulting in fluorescence vanishing. **L** Rational design of the S42M mutation fills the cavity and restores fluorescence. **M** The S42M variant partially rescues fluorescence, but with weak emission.

We also noticed that the phenolic hydroxyl group of DFAME adopts an asymmetric bilateral hydrogen-bonding mode with precise proton donor-acceptor orientations: the O-H moiety of the DFAME phenolic hydroxyl serves as the hydrogen bond proton donor, forming a directional hydrogen bond with the oxygen atom of the buried Tyr (Y90) side chain; meanwhile, the oxygen atom of the same phenolic hydroxyl group acts as the hydrogen bond proton acceptor, engaging with two N-H groups on the inner face of the Arg (49R) guanidinium moiety. The planar, rigid guanidinium group of Arg exhibits a clear dual functional partition: the two inner N-H groups form static, orientation-fixed hydrogen bonds with DFAME, which cooperatively clamp the phenolic hydroxyl group together with Tyr to rigidify the entire fluorophore conformation and suppress intramolec-ular torsion and non-radiative relaxation. Although the remaining outer N-H group of the guanidinium moiety points toward the bulk aqueous solution and forms transient weak hydration interactions, it does not establish a continuous, dynamic hydrogen-bonding network that could mediate energy dissipation. Owing to the high structural rigidity of the guanidinium group, it cannot act as a flexible vibrational conduit to transfer the excited-state energy of DFAME to the bulk solvent. Furthermore, the Tyr residue is fully embedded in the hydrophobic interior of the binding pocket, creating a non-polar, desol-vated microenvironment that blocks solvent reorganization and water-induced quenching pathways. When Arg is mutated to Ala, both the bilateral hydrogen-bonding anchoring and steric pocket sealing effects are completely abolished. Bulk water molecules directly invade the microenvironment of DFAME, triggering intensive solvent-mediated non-radiative relaxation and resulting in the total loss of fluorescence. Overall, all of these mutations resulted in a significant decrease in binding affinity, indicating that the buried hydrogen bonds contribute significantly to the binding affinity and confirming the design accuracy.

We next compared the microenvironment of phenolic hydroxyl in DFAME and natural fluorescent proteins (including GFP and RFP), and found a highly conserved residue preference that His, Thr, Ser, and Asn are the most frequent hydrogen-bonding partners (Fig 4F). Several proteins are displayed in Fig 4G. The results for GFP and RFP, calculated separately, were shown in Fig S8, where they exhibit different preferences. Notably, we identified a distinct mechanism in nucleic acid aptamer systems: in aptamer-fluorophore complexes, K^+^ directly coordinates with the phenolic oxygen of the chromophore (Fig 4G). This cationic interaction stabilizes the electronic cloud and restricts conformational motion through charge screening, representing a unique activation strategy that is clearly different from the hydrogen-bond-dominant mode in both natural fluorescent proteins and our de novo designed protein system. This divergence highlights the adaptive and divergent strategies used by protein versus nucleic acid scaffolds for fluorogenic activation of small-molecule chromophores.

To further dissect the role of the methyl acrylate group of DFAME (opposite the phenolic hydroxyl, Fig 4H), we substituted it with a methyl group to generate a new molecule, i.e. DFHBI (Fig 4I). We found that DFA-5 is unable to activate DFHBI, which may come from the unoccupied space in the pocket (Fig 4J,K). Meanwhile, we performed rational design and introduced a bulky hydrophobic side chain in the protein pocket (S42M, Fig 4L) to backfill the vacated apolar volume. Despite this compensation, the DFA-5-S42M:DFHBI complex presents a reduced contact area, yielding only weak fluorescence activation of DFHBI (Fig 4M and Fig S9).

These results indicate that the methyl acrylate group of DFAME contributes essential van der Waals packing and desolvation-driven stabilization, and extending the conjugated system.

### Rational design of ligands for spectral tuning across the red-to-green range

Building on these observations, we tried to replace the methyl acrylate group with substituted styrene groups or similar groups with five-membered heterocycles. We docked the compound with styrene group (DFST, Fig 5A) into the DFA-5 crystal structure and found that the shared core structure overlays well and preserves the key hydrogen-bond network, while the new phenyl ring snugly fills the apolar subpocket, resulting in a pretty good vina score with -11.9 kcal/mol (Fig 5C). Fluorescence turn-on assay confirmed robust activation with an apparent *K_d_* of 27 nM and exhibits blue-shifted optimal excitation and emission wavelengths 517/582 nm, corresponding to a 65-nm Stokes shift.

**Figure 5:**
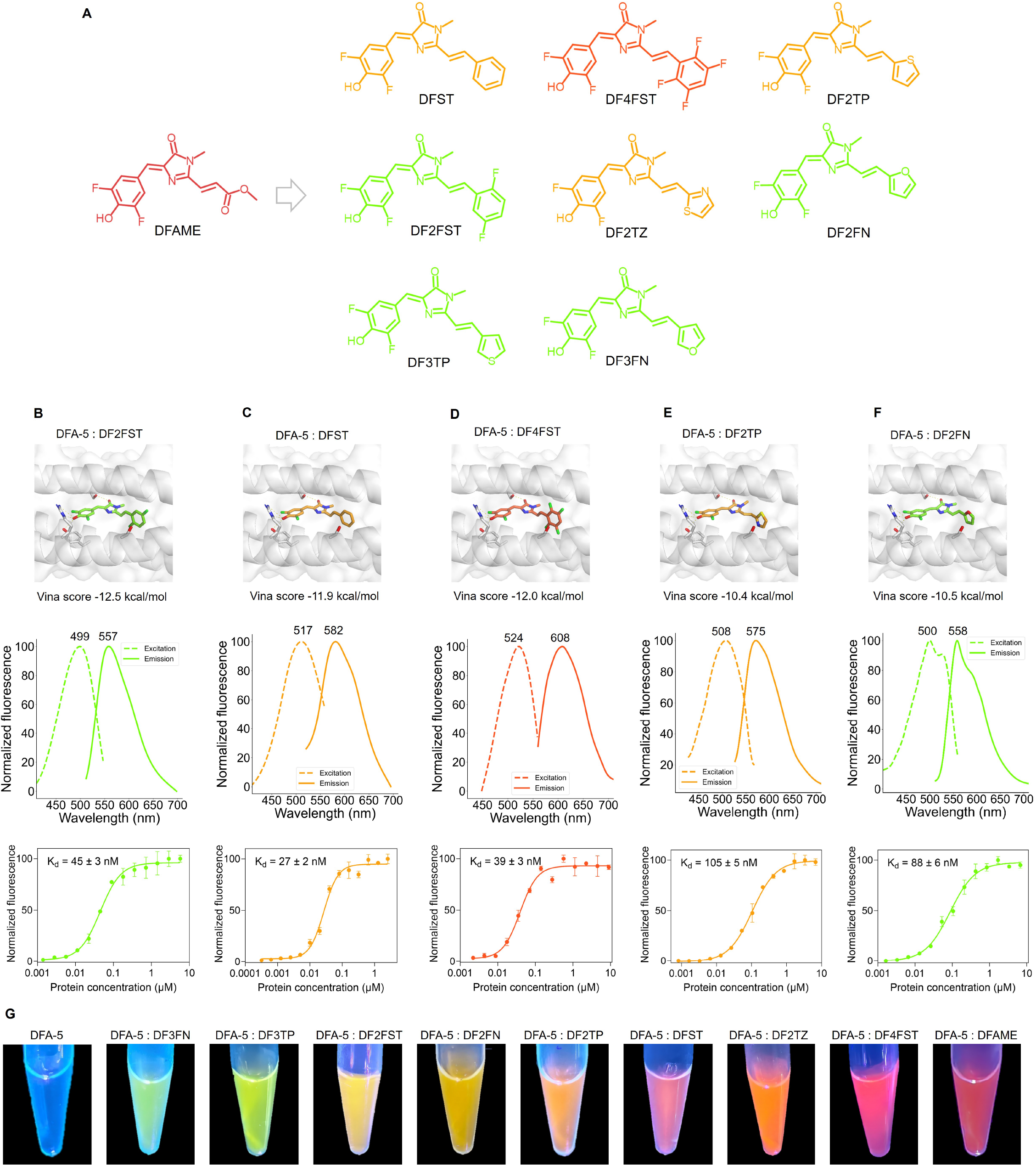
Rational design of small molecule ligands expands the spectral range. **A** Eight molecules designed based on DFAME: DFST, DF2FST, DF4FST, DF2FN, DF3FN, DF2TP, DF3TP and DT2TZ. **B-F** From top to bottom, the three rows show docking poses with affinity scores, wavelength spectra, and binding affinities, respectively. **G** Solutions containing various ligands were illuminated with 365 nm light in the presence of DFA-5.

We further tested the effect of fluorination on the phenyl ring in the styrene group and prepared two compounds: DF2FST and DF4FST (Fig 5A). DF2FST caused spectral blue-shift (499/557 nm) and DF4FST caused red-shift (524/608 nm) compared to DFST while maintaining the similar binding affinity level (Fig 5B,D), which may be attributed to the strong electron-withdrawing effect of the fluorine substituents in DF2FST, leading to a blue shift due to the asymmetric substitution pattern, whereas in DF4FST, the symmetric tetrafluoro substitution and possible hydrogen-bonding interactions with the protein may stabilize the conjugated system and result in a red shift.

We further introduced furan, thiophene, and thiazole groups to replace the phenyl ring in the styrene group, generating DF2FN, DF3FN, DF2TP, DF3TP, and DF2TZ, respectively (Fig 5A). These molecules can also be activated by DFA-5 with mid-nanomolar binding affinities and exhibited excitation wavelengths in the range of 540 to 590 nm (Fig 5E,F and Fig S10). Together, these results indicate that the emission wavelength can be tuned by introducing different substituents on the fluorophore (Fig 5G), thereby shifting the spectral output across the red-to-green spectral range (Table S3).

### Using DFA-5 for cell imaging

As our designed protein DFA-5 exhibits excellent thermostability and broad acid-base tolerance, we further explored its potential for live-cell imaging. Three representative fluorophores, DFAME, DFST and DF2FN, covering red-to-green emission wavelengths, were selected for cellular validation. Upon plasmid transfection with species-optimized codons, both *E. coli* and B16 cells showed obvious fluorescence activation after supplementation with the corresponding exogenous ligands (Fig 6), with signal enhancements of up to 84-fold in *E.coli* and 78-fold in B16 cells. These results demonstrate the robust performance and applicability for live-cell imaging across different cellular systems.

**Figure 6:**
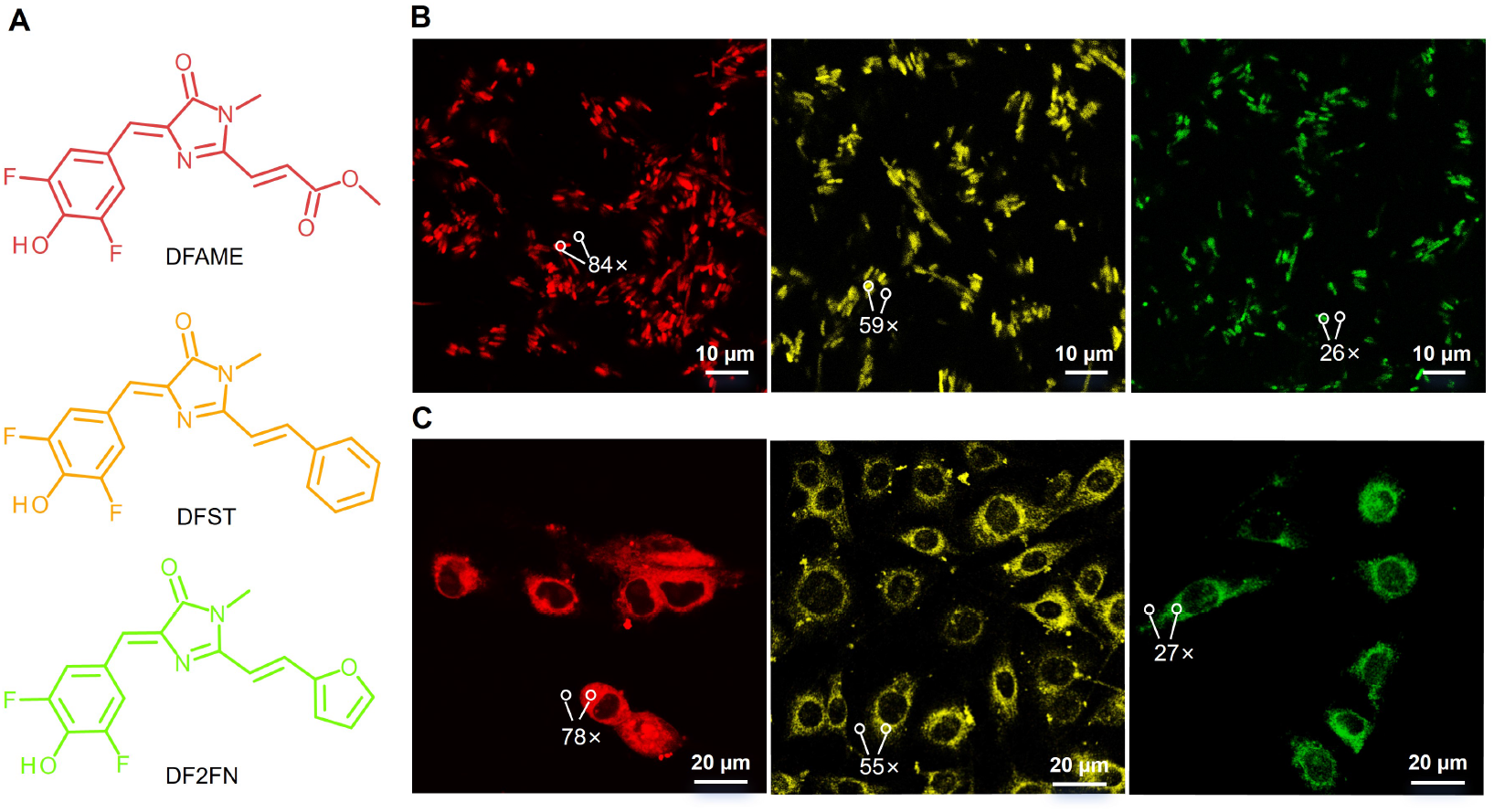
Applications of DFA-5 in cellular imaging. **A** Chemical structures of three representative DFA-5-activated fluorogenic molecules spanning the green-to-red spectral range, together with the application in confocal imaging. **B** and **C** are Confocal images of *E. coli* and B16 cells, repspectively, after treatment adding with the three compounds at 5 µM concentration. The corresponding fluorescence intensities are shown relative to the background signal.

This multi-ligand activation system displays distinctive advantages over conventional fluorescent proteins. First, the compact size of DFA-5, approximately half the molecular weight of canonical GFP/RFP, remarkably reduces cellular expression burden and physiological perturbation. Second, conventional fluorescent proteins generally require several minutes to several hours for the formation and maturation of the intrinsic covalent chromophore [32]. For example, wtGFP and eGFP requires approximately 36 and 15 min, respectively, to reach 50% maturation. In contrast, the fluorescence of DFA-5 is activated rapidly upon ligand binding, making it a suitable probe to detect fast cellular events. Third, conventional imaging usually requires reconstructing distinct plasmids when fusing target proteins with different GFP variants for variable fluorescent emission wavelength, which is time consuming. when DFA-5 is used, only one construct is needed and the fluorescent emission wavelengths can be changed by simply adding different compounds of the DFAME series.

## Discussion

This study establishes a de novo alpha-helical protein environment in which GFP-chromophore-derived ligands can be used to tune fluorogen fluorescence without altering the protein sequence. In conventional GFP systems, spectral properties are primarily diversified by altering the residues surrounding a common chromophore. Here, we instead designed an alpha-helical protein that provides a defined fluorogenic pocket and used a series of GFP-chromophore-derived ligands to vary the optical output while preserving the protein sequence. The resulting system combines a compact and stable protein scaffold with chemically programmable ligand substitution, providing a complementary route to protein-scaffold engineering for the development of small fluorescent probes.

The computational design workflow was essential for obtaining the initial ligand-binding scaffold, while the experimental and structural results also define its current limitations. The experimentally observed ligand conformation was not reproduced in every detail by the predicted complex, particularly in local ligand geometry and interaction orientation. This difference highlights the difficulty of predicting small-molecule binding modes and fluorogenic function from computational models alone. In this context, high-resolution structural analysis is valuable for identifying interaction features that can be used for subsequent ligand and protein optimization. The present results therefore support a combined computational, structural, and experimental strategy for designing functional small-molecule-binding proteins.

A possible explanation for the difficulty of designing efficient helical binders for GFP-chromophore-derived fluorogens is that both binding and fluorescence activation require a precise interaction network. In native GFP-derived environments and recently designed DFHBI-activating beta-barrel proteins [17], the imidazolinone ring nitrogen commonly acts as a hydrogen-bond acceptor and forms a direct polar interaction with the surrounding residues. The geometry and steric constraints of helical scaffolds may make this interaction difficult to reproduce, potentially contributing to the weak binding and limited activation observed in some designs, including those with affinities in the tens-of-micromolar range [33]. Our structure suggests that the successful design may instead stabilize DFAME through an alternative interaction network: its phenolic group is bracketed by the hydroxyl group of Tyr and the guanidinium group of Arg, forming a bilateral polar clamp that restricts ligand motion and maintains a productive orientation. This arrangement may partially compensate for the absence of a canonical hydrogen bond to the imidazolinone nitrogen, suggesting that the efficiency of the present design arises not from reproducing the GFP beta-barrel interaction network, but from replacing a difficult-to-realize contact with distributed interactions that jointly stabilize the ligand and its fluorogenic conformation.

The ligand series demonstrates that a common protein sequence can support substantial spectral diversification through chemical modification of a GFP-chromophore-derived fluorogen. This capability was established here primarily for DFAME-derived ligands, and its transferability to other chromophore families remains to be determined. As a preliminary test of broader scaffold adaptability, structure-guided substitution of three pocket residues also produced a variant capable of binding the chemically distinct drug molecule Benvitimod (Supplementary Results). This result suggests that the scaffold may be retar-geted beyond fluorogenic ligands, although additional measurements and structural basis will be needed to establish the generality of this approach.

Together, these findings support a division of function between the two components of the system: the protein provides a defined microenvironment for fluorogen recognition and fluorescence activation, whereas ligand chemistry provides a route to vary optical output without repeated modification of the protein sequence. The present ligand series shows that this division of function can be implemented for GFP-chromophore-derived fluorogens in a de novo alpha-helical pocket. Further optimization of ligand substituent preferences, pocket interactions, and protein-ligand compatibility may enable coordinated control over emission wavelength, spectral profile, brightness, and binding affinity. The compact size and cellular imaging compatibility of this class of proteins could support applications in live-cell labeling, biosensing, and molecular recognition, while the broader applicability of the design strategy will require validation across additional ligand classes and protein environments.

## Conclusion

In summary, we demonstrate that a compact de novo alpha-helical protein can bind and activate the GFP-chromophore-derived fluorogen DFAME with high affinity. The cocrystal structure revealed the molecular basis of ligand recognition and fluorescence activation, enabling the design of DFAME derivatives that tuned emission from red to green while binding the same protein scaffold without further protein engineering. These findings establish ligand-directed spectral tuning as a complementary strategy to protein-directed chromophore engineering and provide a foundation for extending de novo protein design to other small-molecule ligands.

## Abbreviations

RMSD: root mean square deviation
pLDDT: predicted Local Distance Difference Test
NanoDSF: nano Differential Scanning Fluorimetry
CD: circular dichroism.

## Ethics approval and consent to participate

No applicable.

## Consent for publication

No applicable.

## Availability of data and materials

The scaffolds and source code will be available in https://github.com/PKUliujl/ChameleonBinder.

The co-crystal structure (DFA-5 and DFAME) solved has been deposited in the Protein Data Bank with an accession code of 9XKS.

## Authors’ contributions

JL and LL conceived the project. JL performed the computational studies. YX, CL, JL and YW carried out the experimental work. WZ, JL and ZG collected crystallographic data and solved the crystal structures. ZC synthesized the fluorescent molecules. LL and JL analysed the computational and experimental data. JL wrote the original manuscript with input from all authors. LL and JL revised the manuscript with contributions from all co-authors.

## Acknowledgements

The authors gratefully acknowledge the National Center for Protein Sciences at Peking University in Beijing, China, for the platform of crystallization conditions screening. The authors thank the high-performance computing platform of the Peking-Tsinghua Center for Life Sciences, Peking University, for providing the computational resources. The authors thank the National Facility for Protein Science in Shanghai (NFPS) and Shanghai Synchrotron Radiation Facility for the assistance of crystallographic data collection on the beamline BL10U2.

